# Visual integration of GWAS and differential expression results with the hidecan R package

**DOI:** 10.1101/2023.03.30.535015

**Authors:** Olivia Angelin-Bonnet, Matthieu Vignes, Patrick J. Biggs, Samantha Baldwin, Susan Thomson

## Abstract

**Summary:** We present `hidecan`, an R package for generating visualisations that summarise the results of one or more genome-wide association studies and differential expression analyses, as well as manually curated candidate genes, e.g. extracted from the literature.

**Availability and Implementation:** The hidecan package is implemented in R and is publicly available on the CRAN repository (https://CRAN.R-project.org/package=hidecan) and on GitHub (https://github.com/PlantandFoodResearch/hidecan). A description of the package, as well as a detailed tutorial are available at https://plantandfoodresearch.github.io/hidecan/.

**Supplementary information:** Supplementary data are available.

## 1 Introduction

Genome-wide association studies (GWAS) enable researchers to investigate genomic regions associated with a trait of interest [3]. Often, the ultimate goal of such analyses is to highlight causal genes that are involved in biological processes relating to the phenotype under study; for example to detect disease-related genes that can be targeted by new drugs, or facilitate selection or desirable characteristics in breeding programmes. Therefore, association studies can be complemented with the acquisition of transcriptomics data [15, 17]. These can be used to detect genes whose expression is associated with the trait under study through a differential expression (DE) analysis. The results from both GWAS and DE are typically compared with previous results from the literature, in order to assess which findings are substantiated by previous research, and which are novel results that must be confirmed with follow-up studies.

GWAS results are usually displayed through a Manhattan plot, which represents the score of each genomic variant (i.e. −log10 of its p-value) against its physical position along the genome [14]. DE results are instead typically represented with a volcano plot, which displays the DE score of the genes against their log2-fold change [7]. The drawback of using these plots is that they cannot be easily supplemented with manual information, for example genes of interest from the literature. In addition, Manhattan plots and volcano plots cannot be combined to represent both GWAS and DE results in a single figure. In recent years, the Circos plot has emerged as an attractive design to display information related to genomics data [6], and thus can be used to visualise the results of both GWAS and DE. It is notably possible to combine several layers of information by stacking several circles on top of each other in a single Circos plot [5, 17]. However, the circular layout of Circos plots makes it difficult to precisely compare the different layers and thus to identify genomic regions of interest in which the information from several layers overlap.

We recently proposed a new visualisation, the HIDECAN plot, that integrates GWAS and DE results as well as candidate genes (from the literature) into one graphic [1]. In this paper, we present the corresponding hidecan R package, which can be used to generate HIDECAN plots. The package is publicly available on the CRAN repository (https://CRAN.R-project.org/package=hidecan) and on GitHub (https://github.com/PlantandFoodResearch/hidecan), with a detailed tutorial available at https://plantandfoodresearch.github.io/hidecan/.

## 2 Implementation

The main function of the hidecan package is `the hidecan_plot()` function. It takes as input data-frames of GWAS results, DE results, and candidate gene lists. These data-frames should contain information about the genomic position of the genomic variants and genes (i.e. chromosome and physical position or start and end, in base pairs), as well as GWAS or DE scores and log2-fold change. The user needs to set a significance threshold on the GWAS scores, DE scores and DE log2-fold change in order to select genomics variants and genes that will be displayed in the HIDECAN plot. Several other parameters included in the `hidecan_plot()` function allow the user to control different aspects of the plot, such as which chromosomes should be displayed, or the size of the points in the graphic. The function then returns a HIDECAN plot constructed with the ggplot2 [16], ggrepel [12] and ggnewscale [4] packages (Figure 1 A-D). Supplementary Material 1 details the different steps of the `hidecan_plot()` function to generate a HIDECAN plot from the input datasets.

**Figure 1:**
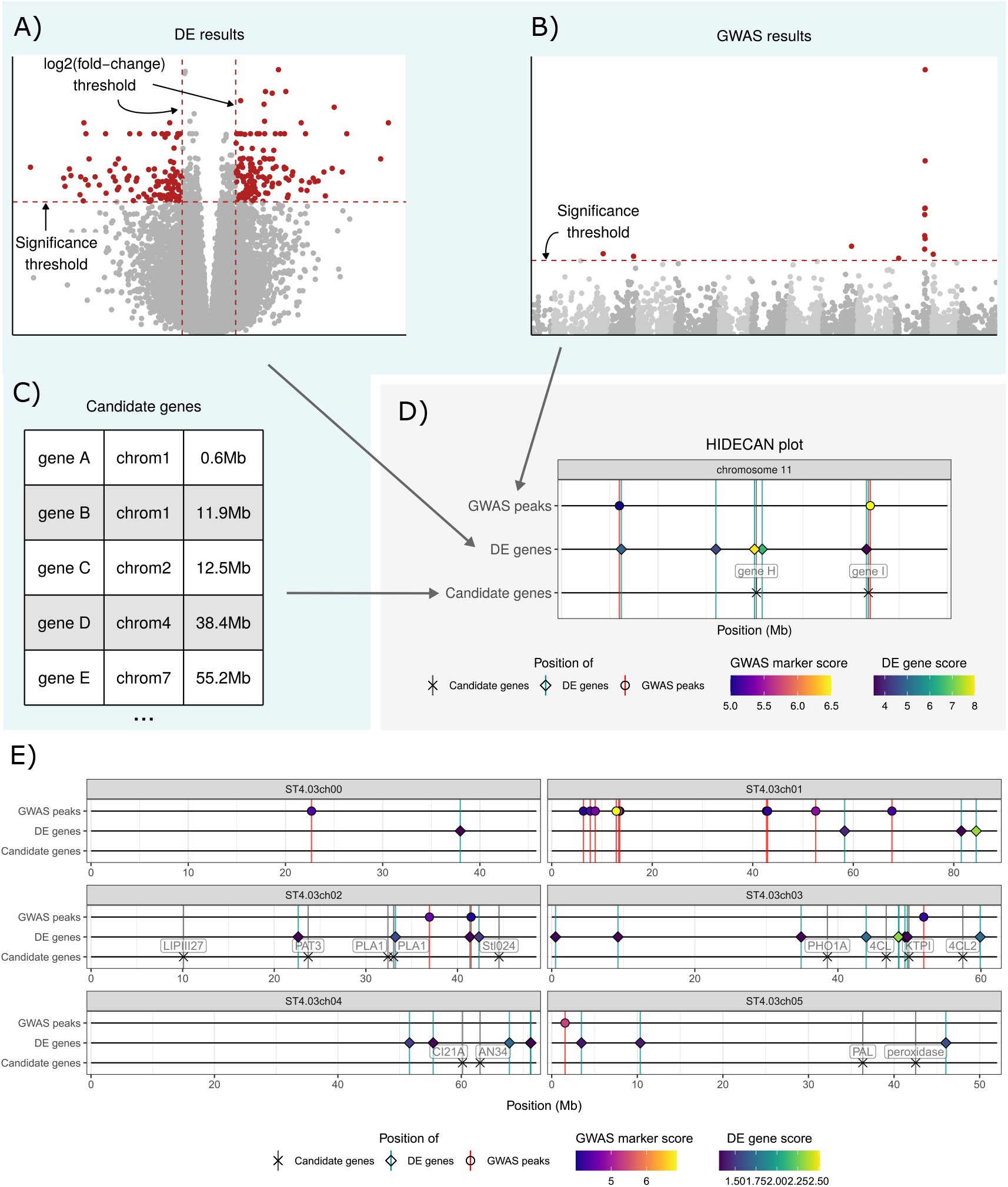
A) to D): Schema of the construction of a HIDECAN plot. The blue box indicates data and plots obtained outside of the hidecan package. A) From a table of differential expression results (represented here with a volcano plot), differentially expressed genes are selected by applying a threshold on the genes’ scores and log2-fold change. B) Similarly, interesting genomic markers are selected from the GWAS results (here represented with a Manhattan plot) by applying a significance threshold on their association score. C) Lastly, a manually curated table of candidate genes of interest can be specified. D) The HIDECAN plot displays the genomic position (x-axis) of the selected genomic variants, differentially expressed genes and candidate genes along each chromosome. The colour of the points represent the GWAS and DE scores. In this example, only chromosome 11 is represented for better clarity. E) A HIDECAN plot generated with the package, from GWAS and DE results on a potato tuber bruising dataset (only chromosomes 0 to 5 are represented for better clarity, see Supplementary Material for the full figure).

Alternatively, a Shiny app is made available, through which the user can construct a HIDECAN plot via a visual interface rather than with code, by providing as input csv files of GWAS results, DE results and candidate genes lists. The generated HIDECAN plot can be saved as either a PNG or PDF file.

Often, the GWAS and DE analyses are performed using popular R packages such as edgeR [10] or DESeq2 [8]. For example, GWASpoly [11] is an R package for performing GWAS for autotetraploid organisms. Through the GWASpoly package, it is possible to obtain GWAS scores for several traits at once and with different genetic models. The package then offers several options to compute a significance threshold on the genomic variants score, for each trait and genetic model. In order to facilitate the construction of HIDECAN plots from GWASpoly results, we are offering a helper function to extract the necessary information directly from the output of the GWASpoly package. The `hidecan_plot_from_ gwaspoly()` function reads in the output from the GWASpoly package and extracts variants scores and significance thresholds to produce a HIDECAN plot. Such helper functions reduce the amount of data wrangling that users must perform to convert the output of these packages into a suitable format for the hidecan package. Due to the modular construction of the package, it is possible to easily implement additional helper functions for other diploid- or polyploid-focused packages if the need arises.

The hidecan package has also been included in the visualisation tool VIEWpoly [13]. VIEWpoly is an R package and Shiny app for visualising and integrating results from polyploid genetic and genomic analysis tools. It provides an intuitive interface that facilitates the interactive exploration of linkage mapping and QTL analysis results, obtained from a suite of packages such as MAPpoly [9] or polyqtlR [2]. Users can explore the QTL results and linkage maps in more detail, or use the implemented genome browser to explore the relationships between QTL regions and a corresponding genome annotation. In the newest version of the package (v0.4.0), it is possible to upload GWAS and DE results as well as candidate gene lists as csv files, or GWASpoly results as RData files, into the VIEWpoly Shiny app. A HIDECAN plot is generated from these input files, using the hidecan package. Users can further explore and customise the HIDECAN plot and save the resulting figure as a file with one of several supported extensions.

## 3 Examples

We demonstrate the use of the hidecan package with a dataset from a breeding population of tetraploid potatoes, originally published in [1]. A GWAS analysis was applied to 72,847 genomic variants obtained from 158 progeny plants from a half-sibling breeding population; the results were compared with candidate genes from previous GWAS or QTL mapping studies on potato tuber bruising. In addition, a DE analysis was used to compare the expression of 25,163 transcribed genes between low- and high-bruising tubers two hours after bruising. A subset of these results was used to generate a HIDECAN plot. Figure 1 E showcases the resulting HIDECAN plot for chromosomes 0 to 5. The full HIDECAN plot, as well as the code used to generate the figure and details about the dataset, are presented in Supplementary Material 2. The data used in this example (i.e. result from the GWAS and DE analyses, as well as the list of candidate genes) is available in the hidecan package through the `get_example_data()` function.

In addition, we showcase how the package can be used to visualise the output from the GWASpoly package without any data manipulation. We use the example dataset from the GWASpoly publication [11], in which genomic information and phenotype measurements are obtained for 221 tetraploid potato lines from the SolCAP diversity panel. We focus on the analysis of tuber eye depth, shape and sucrose content, and perform a GWAS analysis with four different genetic models. The code used for this example and resulting HIDECAN plot are presented in Supplementary Material 2.

## 4 Conclusion

We present the hidecan R package for generating HIDECAN plots that display genomic variants and genes highlighted by GWAS and DE analyses, alongside manually curated candidate genes of interest. The HIDECAN plot provides in a single visualisation a genome-wide overview of these three types of information. Critically, it allows users to easily identify genomic regions in which association with a trait of interest is supported at both the genomics and transcriptomics level, and/or supported by previous findings. The hidecan R package provides simple functions that allow the user to rapidly generate such HIDECAN plots. It also offers a Shiny app which allows users to construct a plot through an interactive web-based interface rather than through code. Finally, it provides helper functions that extract results from the output of packages performing the GWAS or DE analysis, to facilitate the use of the package in an R analysis pipeline.

## 5 Acknowledgements

The authors would like to thank Cristiane Hayumi Taniguti, Oscar Riera-Lizarazu, David Byrne, Marcelo Mollinari from the Tools for Polyploids project for their help and for the opportunity to include the hidecan package into VIEWpoly.

