## Supplementary Material 1 for "Visual integration of GWAS and differential expression results with the hidecan R package"

### Supplementary Material 1 – Package workflow

O. Angelin-Bonnet, M. Vignes, P. J. Biggs, S. Baldwin, S. Thomson

```
library(hidecan)
```

The main function of the `hidecan` package is the `hidecan_plot()` function, which takes as input data-frames of genome-wide association study (GWAS) and differential expression (DE) results, as well as manually curated candidate genes, and returns a HIDECAN plot displaying the genomic position of the significant genomic variants and genes alongside candidate genes. In this document, we present the workflow executed by this function to go from the input datasets to the resulting HIDECAN plot. This workflow is summarised in Figure 1. The following document is also available online at https://plantandfoodresearch.github.io/hidecan/articles/hidecan-step-by-step.html.

#### Input data

The `hidecan` package takes as input tibbles (data-frames) of GWAS and DE results or candidate genes. The input data-frames should contain some mandatory columns, presented below.

A list of example input datasets can be obtained via the `get_example_data()` function:

```
x <- get_example_data()
str(x, max.level = 1)
```

```
List of 3
 $ GWAS: tibble [35,481 x 4] (S3: tbl_df/tbl/data.frame)
 $ DE  : tibble [10,671 x 7] (S3: tbl_df/tbl/data.frame)
 $ CAN : tibble [32 x 6] (S3: tbl_df/tbl/data.frame)
```

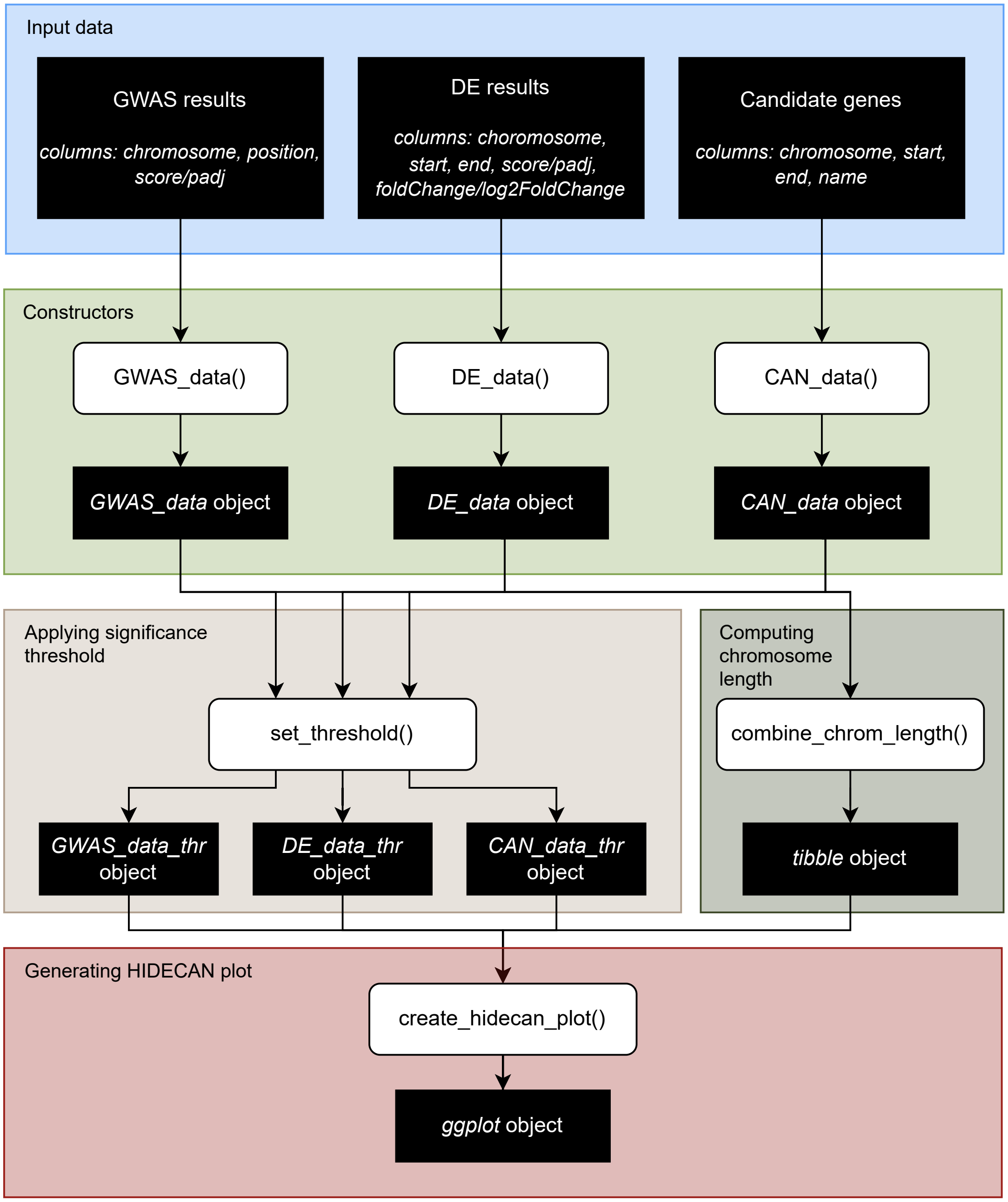


Figure 1: Schema of the `hidecan_plot()` function workflow. Black boxes represent R objects, while white boxes show functions from the `hidecan` package. The different coloured areas correspond to the different steps in the generation of a HIDECAN plot.

#### GWAS results

GWAS results should be provided as a tibble or data-frame, with one row per genetic marker. The data-frame should contain at least the following columns:

- `chromosome`: character column, giving the ID of the chromosome on which each marker is located;
- `position`: numeric column, the physical position along the chromosome (in base pairs - bp) of the marker;
- either `score` or `padj`: numeric column, providing either the score (i.e. `-log10(p-value)`) or the adjusted p-value of the marker. If a `score` column is provided, the `padj` column will be ignored. If only a `padj` column is provided, a `score` column will be constructed as `-log10(padj)`.

Any other column present in the data-frame will be ignored. An example of valid input is shown below:

```
head(x[["GWAS"]])
```

```
# A tibble: 6 x 4
  id                   chromosome position score
  <chr>                <chr>         <dbl> <dbl>
1 ST4.03ch00_45467783 ST4.03ch00 45467783 0.191
2 ST4.03ch01_88589716 ST4.03ch01 88589716 1.84
3 ST4.03ch02_48614228 ST4.03ch02 48614228 0.381
4 ST4.03ch03_62263578 ST4.03ch03 62263578 0.661
5 ST4.03ch04_72139135 ST4.03ch04 72139135 0.640
6 ST4.03ch05_52040302 ST4.03ch05 52040302 0.346
```

#### Differential expression results

DE results should be provided as a tibble or data-frame, with one row per gene. The data-frame should contain at least the following columns:

- `chromosome`: character column, giving the ID of the chromosome on which each gene is located;
- `start` and `end`: numeric columns, giving the starting and end position of the gene along the chromosome (in bp). These two columns will be used to calculate the position of the gene as the half-way point between the start and end of the gene.

- either `score` or `padj`: numeric column, providing either the score (i.e. `-log10(p-value)`) or the adjusted p-value of the gene. If a `score` column is provided, the `padj` column will be ignored. If only a `padj` column is provided, a `score` column will be constructed as `-log10(padj)`.
- either `foldChange` or `log2FoldChange`: numeric column, giving either the fold-change or log2(fold-change) of the gene. If a `log2FoldChange` column is provided, the `foldChange` column will be ignored. If only a `foldChange` column is provided, a `log2FoldChange` will be constructed as `log2(foldChange)`.

Any other column present in the data-frame will be ignored. An example of valid input is shown below:

```
head(x[["DE"]])
```

```
# A tibble: 6 x 7
  gene                 chromosome   padj log2FoldChange    start      end label
  <chr>                <chr>       <dbl>          <dbl>    <dbl>    <dbl> <chr>
1 PGSC0003DMG400032056 ST4.03ch00  0.787        0.0114  45813195 45813526 Prote~
2 PGSC0003DMG400018039 ST4.03ch01  0.630        0.00529 88623473 88627702 PhD-f~
3 PGSC0003DMG400020231 ST4.03ch02  0.864        0.00362 48563271 48578978 Acety~
4 PGSC0003DMG400009197 ST4.03ch03  0.530        0.0320  62256322 62258929 Phosp~
5 PGSC0003DMG403025662 ST4.03ch04  0.975        0.00225 72168842 72170119 Conse~
6 PGSC0003DMG400023316 ST4.03ch05 NA          -0.000726 52039916 52040326 Conse~
```

(Note that in the example dataset, some genes have missing values in the `padj` column; this corresponds to genes that have been filtered out via independent filtering in the DESeq2 package).

#### Candidate genes

A list of candidate genes (e.g. genes previously found associated with a trait of interest based on literature search) can be provided as a tibble or data-frame, with one row per gene. This data-frame can also contain variants of interest (see below). The data-frame should contain at least the following columns:

- `chromosome`: character column, giving the ID of the chromosome on which each gene is located;
- `start` and `end`: numeric columns, giving the starting and end position of the gene along the chromosome (in bp). These two columns will be used to calculate the position of the gene as the half-way point between the start and end of the gene. For genomic variants

or markers, simply set both the `start` and `end` columns to the physical position of the marker.

- `name`: character column, giving the name of the candidate gene that will be displayed in the HIDECAN plot. Set to `NA` for any subset of genes to remove their label in the plot (can help if many genes are very close, to avoid cluttering the plot).

Any other column present in the data-frame will be ignored. An example of valid input is shown below:

```r
head(x[["CAN"]])
```

```
# A tibble: 6 x 6
  id                   chromosome    start      end name       gene_name
  <chr>                <chr>         <dbl>    <dbl> <chr>      <chr>
1 PGSC0003DMG400003155 ST4.03ch03 46757152 46762127 4CL        4-coumarate-CoA ~
2 PGSC0003DMG400014223 ST4.03ch03 57466692 57469946 4CL2       4-coumarate-CoA ~
3 PGSC0003DMG400011189 ST4.03ch07  1001854  1006278 HQT        HQT
4 PGSC0003DMG400005492 ST4.03ch05 36342746 36347409 PAL        phenylalanine am~
5 PGSC0003DMG400005279 ST4.03ch05 42523943 42525912 peroxidase peroxidase
6 PGSC0003DMG400007782 ST4.03ch03 38537202 38540209 PHO1A      PHO1A
```

#### Constructors – formatting the input data

Under the hood, the `hidecan` package relies on S3 classes, which are really just tibbles with specific columns. The constructors for these S3 classes perform a series of checks and computations to make sure that all of the required columns (such as chromosome, score, position) are present in the data.

- GWAS results data-frames are turned into `GWAS_data` objects through the `GWAS_data()` constructor:

```r
gwas_data <- GWAS_data(x[["GWAS"]])
class(gwas_data)
```

```
[1] "GWAS_data"  "tbl_df"     "tbl"        "data.frame"
```

```r
head(gwas_data)
```

```
# A tibble: 6 x 4
  id                   chromosome position score
  <chr>                <chr>         <dbl> <dbl>
1 ST4.03ch00_45467783 ST4.03ch00 45467783 0.191
2 ST4.03ch01_88589716 ST4.03ch01 88589716 1.84
3 ST4.03ch02_48614228 ST4.03ch02 48614228 0.381
4 ST4.03ch03_62263578 ST4.03ch03 62263578 0.661
5 ST4.03ch04_72139135 ST4.03ch04 72139135 0.640
6 ST4.03ch05_52040302 ST4.03ch05 52040302 0.346
```

- DE results data-frames are turned into `DE_data` objects through the `DE_data()` constructor:

```
de_data <- DE_data(x[["DE"]])
class(de_data)
```

```
[1] "DE_data"    "tbl_df"     "tbl"        "data.frame"
```

```
head(de_data)
```

```
# A tibble: 6 x 9
  gene                 chrom~1   padj log2Fo~2  start    end label   score posit~3
  <chr>                <chr>    <dbl>    <dbl>  <dbl>  <dbl> <chr>   <dbl>   <dbl>
1 PGSC0003DMG400032~ ST4.03~  0.787  1.14e-2 4.58e7 4.58e7 Prot~  0.104   4.58e7
2 PGSC0003DMG400018~ ST4.03~  0.630  5.29e-3 8.86e7 8.86e7 PhD--~  0.201   8.86e7
3 PGSC0003DMG400020~ ST4.03~  0.864  3.62e-3 4.86e7 4.86e7 Acet~  0.0637  4.86e7
4 PGSC0003DMG400009~ ST4.03~  0.530  3.20e-2 6.23e7 6.23e7 Phos~  0.276   6.23e7
5 PGSC0003DMG403025~ ST4.03~  0.975  2.25e-3 7.22e7 7.22e7 Cons~  0.0109  7.22e7
6 PGSC0003DMG400023~ ST4.03~ NA     -7.26e-4 5.20e7 5.20e7 Cons~ NA       5.20e7
# ... with abbreviated variable names 1: chromosome, 2: log2FoldChange,
#   3: position
```

- Candidate genes data-frames are turned into `CAN_data` objects through the `CAN_data()` constructor:

```
## CAN_data constructor
can_data <- CAN_data(x[["CAN"]])
class(can_data)
```

```
[1] "CAN_data"   "tbl_df"     "tbl"        "data.frame"
```

```r
head(can_data)
```

```
# A tibble: 6 x 7
  id                   chromosome    start      end name       gene_name posit~1
  <chr>                <chr>         <dbl>    <dbl> <chr>      <chr>       <dbl>
1 PGSC0003DMG400003155 ST4.03ch03 46757152 46762127 4CL        4-coumar~  4.68e7
2 PGSC0003DMG400014223 ST4.03ch03 57466692 57469946 4CL2       4-coumar~  5.75e7
3 PGSC0003DMG400011189 ST4.03ch07  1001854  1006278 HQT        HQT        1.00e6
4 PGSC0003DMG400005492 ST4.03ch05 36342746 36347409 PAL        phenylal~  3.63e7
5 PGSC0003DMG400005279 ST4.03ch05 42523943 42525912 peroxidase peroxida~  4.25e7
6 PGSC0003DMG400007782 ST4.03ch03 38537202 38540209 PHO1A      PHO1A      3.85e7
# ... with abbreviated variable name 1: position
```

These constructors will throw an error if a required column is missing from the input data (e.g. no chromosome column):

```r
library(dplyr)

gwas_wrong_input <- x[["GWAS"]] |>
  select(-chromosome)
GWAS_data(gwas_wrong_input)
```

```
Error: Input data-frame is missing the following columns: 'chromosome'.
```

They will also compute marker or gene scores from adjusted p-values if necessary. For example, for DE results, if we provide a `padj` column (with the adjusted p-values of the genes) rather than a `score` column, the constructor will compute the `score` column based on the `padj` column. You can also notice that a `position` column is computed based on the start and end of the genes:

```r
## Input tibble
head(x[["DE"]])
```

```
# A tibble: 6 x 7
  gene                 chromosome   padj log2FoldChange    start      end label
  <chr>                <chr>       <dbl>          <dbl>    <dbl>    <dbl> <chr>
1 PGSC0003DMG400032056 ST4.03ch00  0.787        0.0114   45813195 45813526 Prote~
2 PGSC0003DMG400018039 ST4.03ch01  0.630        0.00529  88623473 88627702 PhD-f~
3 PGSC0003DMG400020231 ST4.03ch02  0.864        0.00362  48563271 48578978 Acety~
```

```
4 PGSC0003DMG400009197 ST4.03ch03  0.530       0.0320   62256322 62258929 Phosp~
5 PGSC0003DMG403025662 ST4.03ch04  0.975       0.00225  72168842 72170119 Conse~
6 PGSC0003DMG400023316 ST4.03ch05 NA          -0.000726 52039916 52040326 Conse~
```

```
## Output of the DE_data constructor
head(de_data)
```

```
# A tibble: 6 x 9
  gene                 chrom~1   padj log2Fo~2  start    end label   score posit~3
  <chr>                <chr>    <dbl>    <dbl>  <dbl>  <dbl> <chr>   <dbl>   <dbl>
1 PGSC0003DMG400032~ ST4.03~  0.787  1.14e-2 4.58e7 4.58e7 Prot~  0.104   4.58e7
2 PGSC0003DMG400018~ ST4.03~  0.630  5.29e-3 8.86e7 8.86e7 PhD--  0.201   8.86e7
3 PGSC0003DMG400020~ ST4.03~  0.864  3.62e-3 4.86e7 4.86e7 Acet~  0.0637  4.86e7
4 PGSC0003DMG400009~ ST4.03~  0.530  3.20e-2 6.23e7 6.23e7 Phos~  0.276   6.23e7
5 PGSC0003DMG403025~ ST4.03~  0.975  2.25e-3 7.22e7 7.22e7 Cons~  0.0109  7.22e7
6 PGSC0003DMG400023~ ST4.03~ NA     -7.26e-4 5.20e7 5.20e7 Cons~ NA       5.20e7
# ... with abbreviated variable names 1: chromosome, 2: log2FoldChange,
#   3: position
```

#### Computing chromosome length

Once the input datasets have been formatted appropriately, they are used to compute the length of the chromosomes present in the data. This is done through the `combine_chrom_length()` function, which is applied to a list of `GWAS_data`, `DE_data` or `CAN_data` objects:

```
chrom_length <- combine_chrom_length(list(gwas_data,
                                          de_data,
                                          can_data))
chrom_length
```

```
# A tibble: 13 x 2
   chromosome   length
   <chr>         <dbl>
 1 ST4.03ch00 45813526
 2 ST4.03ch01 88627702
 3 ST4.03ch02 48614228
 4 ST4.03ch03 62263578
 5 ST4.03ch04 72170119
 6 ST4.03ch05 52040326
```

```
 7 ST4.03ch06 59476545
 8 ST4.03ch07 56715111
 9 ST4.03ch08 56937627
10 ST4.03ch09 61539681
11 ST4.03ch10 59687482
12 ST4.03ch11 45409456
13 ST4.03ch12 61152223
```

The function works by calling for each element in the list the `compute_chrom_length()` function. The function, according to whether the input is a tibble of markers (`GWAS_data`) or genes (`DE_data` or `CAN_data`), looks for the maximum value in either the `position` column (for markers) or the `end` column (for genes).

```
head(compute_chrom_length(gwas_data), 3)
```

```
# A tibble: 3 x 2
  chromosome   length
  <chr>         <dbl>
1 ST4.03ch00 45467783
2 ST4.03ch01 88589716
3 ST4.03ch02 48614228
```

```
head(compute_chrom_length(de_data), 3)
```

```
# A tibble: 3 x 2
  chromosome   length
  <chr>         <dbl>
1 ST4.03ch00 45813526
2 ST4.03ch01 88627702
3 ST4.03ch02 48578978
```

#### Applying threshold

Next, the GWAS and DE results tibbles are filtered according to a threshold, in order to retain the significant markers or genes. This is done through the `apply_threshold()` function. This function has two (rather self-explanatory) arguments: `score_thr` and `log2fc_thr`.

When applied to a `GWAS_data` object, the function filters markers with a score above the value set with `score_thr` argument (the `log2fc_thr` argument is ignored), and returns an object of class `GWAS_data_thr`:

```r
dim(gwas_data)
```

```
[1] 35481     4
```

```r
gwas_data_thr <- apply_threshold(gwas_data,
                                 score_thr = 4)
class(gwas_data_thr)
```

```
[1] "GWAS_data_thr" "tbl_df"        "tbl"           "data.frame"
```

```r
dim(gwas_data_thr)
```

```
[1] 37  4
```

```r
head(gwas_data_thr)
```

```
# A tibble: 6 x 4
  id                   chromosome position score
  <chr>                <chr>         <dbl> <dbl>
1 ST4.03ch00_22680252 ST4.03ch00 22680252  4.41
2 ST4.03ch01_6317643  ST4.03ch01  6317643  4.15
3 ST4.03ch01_7671100  ST4.03ch01  7671100  4.43
4 ST4.03ch01_8653747  ST4.03ch01  8653747  4.69
5 ST4.03ch01_12842648 ST4.03ch01 12842648  6.85
6 ST4.03ch01_13334335 ST4.03ch01 13334335  5.24
```

For a `DE_data` object, the `apply_threshold()` function filters genes based on both their score and log2(fold-change), and returns an object of class `DE_data_thr`:

```r
dim(de_data)
```

```
[1] 10671     9
```

```r
de_data_thr <- apply_threshold(de_data,
                               score_thr = 2,
```

```r
                              log2fc_thr = 0.5)
class(de_data_thr)
```

```
[1] "DE_data_thr" "tbl_df"      "tbl"         "data.frame"
```

```r
dim(de_data_thr)
```

```
[1] 2 9
```

```r
head(de_data_thr)
```

```
# A tibble: 2 x 9
  gene                 chrom~1    padj log2F~2  start    end label score posit~3
  <chr>                <chr>     <dbl>   <dbl>  <dbl>  <dbl> <chr> <dbl>   <dbl>
1 PGSC0003DMG400015948 ST4.03~ 0.00480   0.757 3.30e7 3.30e7 Cons~  2.32  3.30e7
2 PGSC0003DMG400029856 ST4.03~ 0.00305   0.691 5.82e7 5.82e7 N-ac~  2.52  5.82e7
# ... with abbreviated variable names 1: chromosome, 2: log2FoldChange,
#   3: position
```

Finally, if applied to a CAN_data object, the `apply_threshold()` function simply returns the input tibble as an object of class `CAN_data_thr`:

```r
dim(can_data)
```

```
[1] 32  7
```

```r
can_data_thr <- apply_threshold(can_data,
                               score_thr = 2,
                               log2fc_thr = 0.5)
class(can_data_thr)
```

```
[1] "CAN_data_thr" "tbl_df"       "tbl"          "data.frame"
```

```r
dim(can_data_thr)
```

```
[1] 32  7
```

```r
head(can_data_thr)
```

```
# A tibble: 6 x 7
  id                   chromosome    start      end name       gene_name posit~1
  <chr>                <chr>         <dbl>    <dbl> <chr>      <chr>       <dbl>
1 PGSC0003DMG400003155 ST4.03ch03 46757152 46762127 4CL        4-coumar~  4.68e7
2 PGSC0003DMG400014223 ST4.03ch03 57466692 57469946 4CL2       4-coumar~  5.75e7
3 PGSC0003DMG400011189 ST4.03ch07  1001854  1006278 HQT        HQT        1.00e6
4 PGSC0003DMG400005492 ST4.03ch05 36342746 36347409 PAL        phenylal~  3.63e7
5 PGSC0003DMG400005279 ST4.03ch05 42523943 42525912 peroxidase peroxida~  4.25e7
6 PGSC0003DMG400007782 ST4.03ch03 38537202 38540209 PHO1A      PHO1A      3.85e7
# ... with abbreviated variable name 1: position
```

As for the `GWAS_data`, `DE_data` or `CAN_data` objects, the `GWAS_data_thr`, `DE_data_thr` and `CAN_data_thr` objects are really just tibbles.

### Creating the HIDECAN plot

Finally, the filtered datasets are combined into a list and passed on to the `create_hidecan_plot()` function, along with the tibble of chromosome length. This is the function that generates the HIDECAN `ggplot`:

```r
create_hidecan_plot(
  list(gwas_data_thr,
       de_data_thr,
       can_data_thr),
  chrom_length
)
```

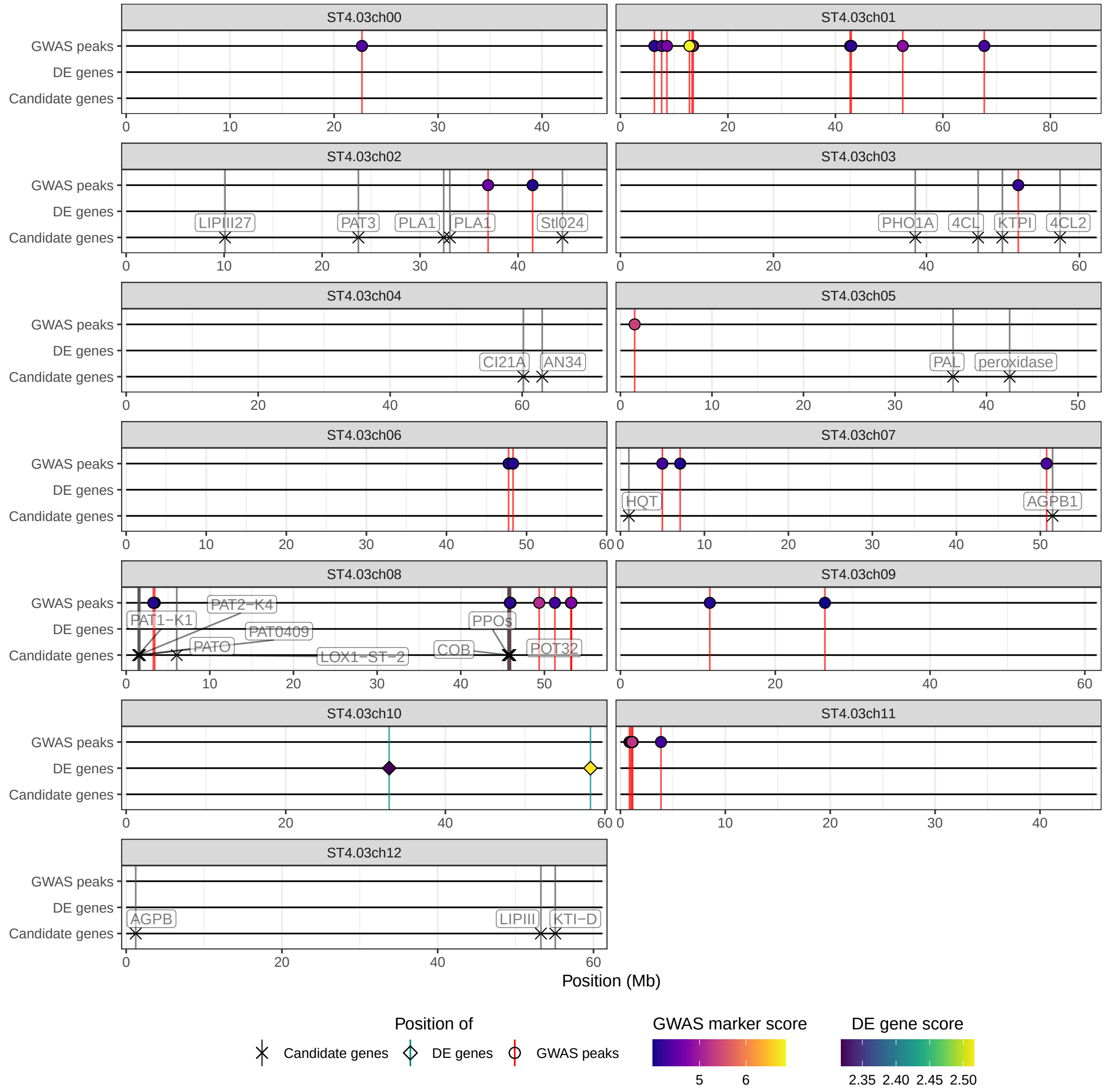


This function shares most of its arguments with the `hidecan_plot()` wrapper for controlling different aspects of the plot.
