## Supplementary Material 2 for "Visual integration of GWAS and differential expression results with the hidecan R package"

**Supplementary Material 2 – Examples**

O. Angelin-Bonnet, M. Vignes, P. J. Biggs, S. Baldwin, S. Thomson

```r
library(hidecan)
```

### Example 1 – potato tuber bruising

#### The data

The dataset used in this example is presented in (Angelin-Bonnet et al. 2023). In this study, tetraploid potato plants from a half-sibling breeding population were used to assess the genetic components of tuber bruising. Capture sequencing was used to obtain genomic information about the individuals, and a genome-wide association study (GWAS) was performed on 72,847 genomic biallelic variants obtained from 158 plants for which a bruising score was measured. The GWAS analysis was carried with the `GWASpoly` package.

In addition, expression data was obtained for 25,163 transcribed genes, and a differential expression (DE) analysis was carried out between 41 low- and 33 high-bruising samples.

Finally, a literature search yielded a list of 42 candidate genes identified in previous studies as involved in potato tuber bruising mechanisms.

A subset of the GWAS and DE results, as well as the list of candidate genes from the literature, are made available in the `hidecan` package through the `get_example_data()` function. From the complete GWAS results table, half of the genomic variants with a GWAS score $< 3.5$ were randomly selected and consequently discarded, yielding a dataset with GWAS scores for 35,481 variants. Similarly, half of the transcribed genes in the DE results table with an adjusted p-value $> 0.05$ were randomly selected and discarded, yielding a dataset with DE results for 10,671 transcribed genes. This filtering was performed to reduce the size of the datasets (in accordance with CRAN policies), but ensures that all significant markers and genes are retained in the datasets. Finally, some of the candidate genes located on chromosome 3 were removed from the example dataset for better clarity in the resulting HIDECAN plot, leaving 32 candidate genes.

The example data can be obtained with:

```
data <- get_example_data()

str(data, max.level = 1)
```

```
List of 3
 $ GWAS:Classes 'tbl_df', 'tbl' and 'data.frame':   35481 obs. of  4 variables:
 $ DE  :Classes 'tbl_df', 'tbl' and 'data.frame':   10671 obs. of  7 variables:
 $ CAN :Classes 'tbl_df', 'tbl' and 'data.frame':   32 obs. of  6 variables:
```

The `get_example_data()` function returns a list of 3 data-frames: the GWAS results table (`GWAS` element), the DE results table (`DE` element) and the candidate genes table (`CAN` element).

The GWAS results table contains, for each genomic variant:

- its genomic position (chromosome and physical position in bp) on the potato genome (PGSC-DM v4.03);
- its GWAS score (i.e. $-\log_{10}$(p-value)).

```
head(data[["GWAS"]])
```

```
                   id chromosome position     score
1 ST4.03ch00_45467783 ST4.03ch00 45467783 0.1912643
2 ST4.03ch01_88589716 ST4.03ch01 88589716 1.8353132
3 ST4.03ch02_48614228 ST4.03ch02 48614228 0.3807299
4 ST4.03ch03_62263578 ST4.03ch03 62263578 0.6609841
5 ST4.03ch04_72139135 ST4.03ch04 72139135 0.6396985
6 ST4.03ch05_52040302 ST4.03ch05 52040302 0.3463201
```

The DE results tables contains, for each transcribed gene:

- its genomic position (i.e. chromosome, as well as start and end positions in bp);
- its adjusted p-value and log2-fold change from the differential expression analysis;
- a label describing the function of the gene (ignored by `hidecan`).

```r
head(data[["DE"]])
```

```
                 gene chromosome      padj log2FoldChange    start      end
1 PGSC0003DMG400032056 ST4.03ch00 0.7874256   0.0114310427 45813195 45813526
2 PGSC0003DMG400018039 ST4.03ch01 0.6300701   0.0052887972 88623473 88627702
3 PGSC0003DMG400020231 ST4.03ch02 0.8636122   0.0036185375 48563271 48578978
4 PGSC0003DMG400009197 ST4.03ch03 0.5301977   0.0319719569 62256322 62258929
5 PGSC0003DMG403025662 ST4.03ch04 0.9752212   0.0022488677 72168842 72170119
6 PGSC0003DMG400023316 ST4.03ch05        NA  -0.0007257173 52039916 52040326
                              label
1               Protein transporter
2                 PhD-finger protein
3              Acetyl-CoA synthetase
4 Phosphatidylcholine transfer protein
5   Conserved gene of unknown function
6   Conserved gene of unknown function
```

The candidate gene table contains, for each gene extracted from the literature:

- its genomic position (i.e. chromosome, as well as start and end positions in bp);
- its label, corresponding to a short version of the complete gene name;
- its complete gene name (ignored by `hidecan`).

```r
head(data[["CAN"]])
```

```
                    id chromosome    start      end       name
1 PGSC0003DMG400003155 ST4.03ch03 46757152 46762127        4CL
2 PGSC0003DMG400014223 ST4.03ch03 57466692 57469946       4CL2
3 PGSC0003DMG400011189 ST4.03ch07  1001854  1006278        HQT
4 PGSC0003DMG400005492 ST4.03ch05 36342746 36347409        PAL
5 PGSC0003DMG400005279 ST4.03ch05 42523943 42525912 peroxidase
6 PGSC0003DMG400007782 ST4.03ch03 38537202 38540209      PHO1A
                  gene_name
1     4-coumarate-CoA ligase
2   4-coumarate-CoA ligase 2
3                        HQT
4 phenylalanine ammonia-lyase
5                 peroxidase
6                      PHO1A
```

#### Constructing the HIDECAN plot

From these three tables, a HIDECAN plot can be generated through the `hidecan_plot()` function. This requires to set a significance threshold on the GWAS score of the genomic variants, as well as on the DE score and log2-fold change of the transcribed genes, which will determine which variants and genes are represented in the plot. For this example, we will set the significance threshold for the GWAS analysis to 4 (corresponding to a p-value of $1 \times 10^{-4}$). We will use for the DE results a significance threshold of 1.3 (corresponding to an adjusted p-value of 0.05) and a log2 fold-change threshold of 0 (which amounts to no filtering based on the log2-fold change). These thresholds are set through the `score_thr_gwas`, `score_thr_de` and `log2fc_thr` arguments, respectively.

```
hidecan_plot(
  gwas_list = data[["GWAS"]], ## data-frame of GWAS results
  de_list = data[["DE"]],     ## data-frame of DE results
  can_list = data[["CAN"]],   ## data-frame of candidate genes
  score_thr_gwas = 4,         ## sign. threshold for GWAS
  score_thr_de = 1.3,         ## sign. threshold for DE
  log2fc_thr = 0              ## log2-fold change threshold for DE
)
```

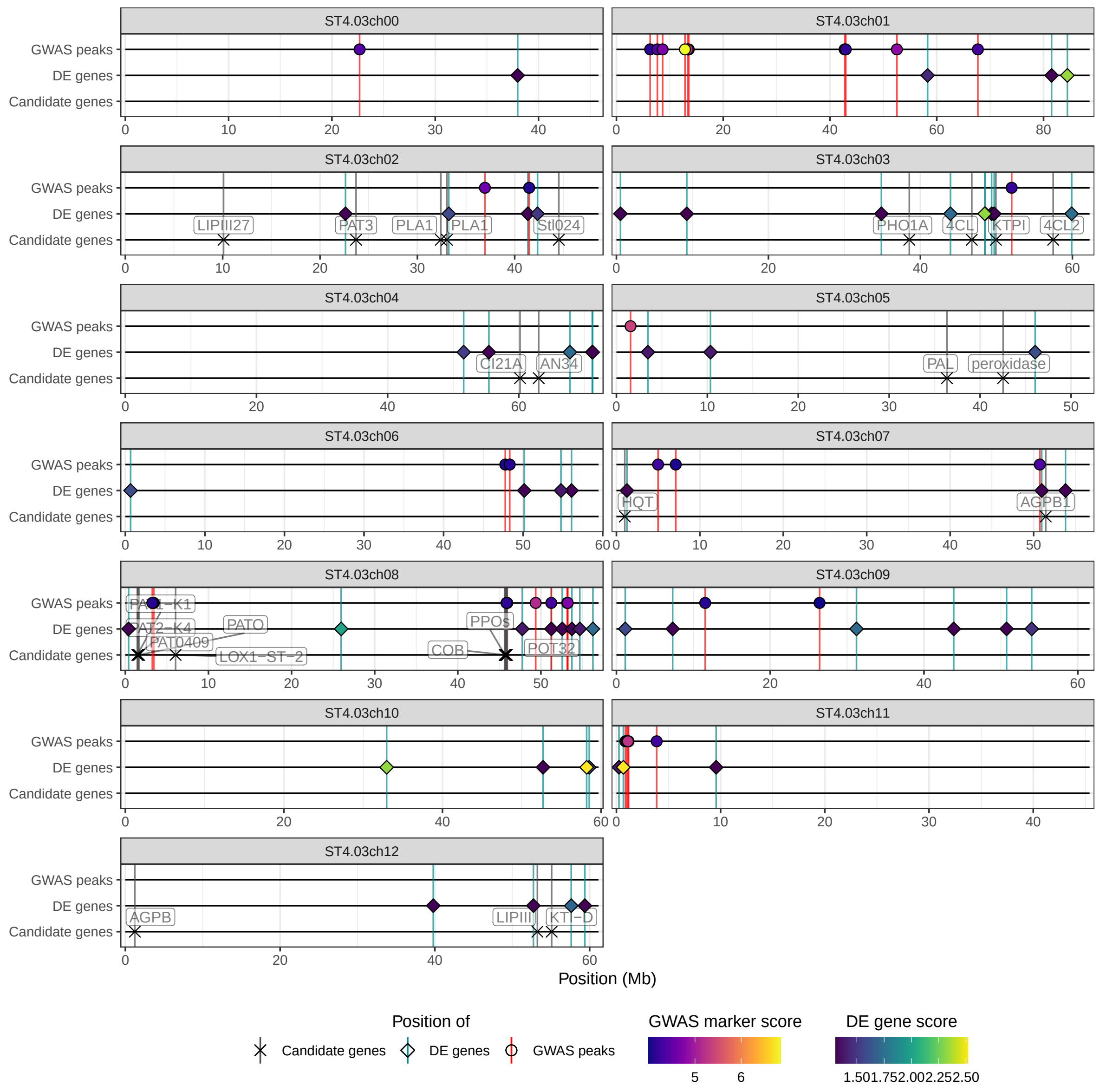


Note that, when executing the following code in an RMarkdown or Quarto document, the resulting plot is too large for the default figure dimensions, thus resulting in the following error when rendering the document:

```
Error in grid.Call(C_convert, x, as.integer(whatfrom), as.integer(whatto),  :
  Viewport has zero dimension(s)
```

In order to avoid this error, the chunk options `fig.width` and `fig.height` are set to a higher value (here 10).

This example, as well as a demonstration of additional options available through the `hidecan_plot()` function are available online at https://plantandfoodresearch.github.io/hidecan/articles/hidecan.html.

### Example 2 – GWASpoly output

#### The data

In this second example, we demonstrate how the output of the `GWASpoly` package can be used directly to generate a HIDECAN plot without formatting of the data. A tutorial for the `GWASpoly` package can be found at https://jendelman.github.io/GWASpoly/GWASpoly.html.

This example is based on the example dataset provided in the publication associated with the `GWASpoly` package (Rosyara et al. 2016), containing genomic and phenotypic information for 221 tetraploid potato lines from the SolCAP diversity panel. In this example, we focus on three of the recorded traits: tuber eye depth, tuber shape, and tuber sucrose content. We will test out four different genetic models for the genotype-phenotype association model: general, additive, simplex reference-dominant and simplex alternate-dominant. For details about the different genetic models, see the GWASpoly publication (Rosyara et al. 2016).

The GWAS analysis can be run with the `GWASpoly` package with the following code:

```
library(GWASpoly)

## Path to genomic and phenotype data files
genofile <- system.file("extdata", "TableS1.csv", package = "GWASpoly")
phenofile <- system.file("extdata", "TableS2.csv", package = "GWASpoly")

## Reading SolCAP data
data <- read.GWASpoly(
  ploidy = 4,
  pheno.file = phenofile,
  geno.file = genofile,
  format = "ACGT",
  n.traits = 13,
  delim = ","
)
```

```
Number of polymorphic markers: 3521
Missing marker data imputed with population mode
```

```
N = 187 individuals with phenotypic and genotypic information
Detected following fixed effects:
Grp1
Grp2
Grp3
Grp4
Detected following traits:
total_yield
chip_color
tuber_eye_depth
tuber_shape
tuber_size
tuber_length
tuber_width
sucrose
log10_glucose
log10_fructose
malic_acid
vine_maturity_95d
vine_maturity_120d
```

```r
## Computing K matrix
data.original <- set.K(
  data,
  LOCO = FALSE,
  n.core = 2
)

## Performing GWAS
gwaspoly_res <- GWASpoly(
  data.original,
  models = c("general", "additive", "1-dom"),
  traits = c("tuber_eye_depth", "tuber_shape", "sucrose"),
  n.core = 2
)
```

```
Using default value for max.geno.freq = 1 - 5/N
Analyzing trait: tuber_eye_depth
P3D approach: Estimating variance components...Completed
Testing markers for model: general
Testing markers for model: additive
```

```
Testing markers for model: 1-dom-alt
Testing markers for model: 1-dom-ref
Analyzing trait: tuber_shape
P3D approach: Estimating variance components...Completed
Testing markers for model: general
Testing markers for model: additive
Testing markers for model: 1-dom-alt
Testing markers for model: 1-dom-ref
Analyzing trait: sucrose
P3D approach: Estimating variance components...Completed
Testing markers for model: general
Testing markers for model: additive
Testing markers for model: 1-dom-alt
Testing markers for model: 1-dom-ref
```

```
## Computing significance threshold
## Object returned by get_gwaspoly_example_data()
gwaspoly_res_thr <- set.threshold(
  gwaspoly_res,
  method = "M.eff",
  level = 0.05
)
```

```
Warning: as(<matrix>, "dspMatrix") is deprecated since Matrix 1.5-0; do
as(as(as(., "dMatrix"), "symmetricMatrix"), "packedMatrix") instead
```

```
Thresholds
                general additive 1-dom-alt 1-dom-ref
tuber_eye_depth    4.67     4.67      4.46      4.49
tuber_shape        4.67     4.67      4.46      4.49
sucrose            4.67     4.67      4.46      4.49
```

The resulting `gwaspoly_res_thr` object contains information about the genomic variants as well as their score for each of the traits and genetic models tested, and the significance threshold for each trait and genetic model used. As an alternative to running the analysis through the code presented above, the `gwaspoly_res_thr` object is available through the `hidecan` package, and can be loaded with the following command:

```
gwaspoly_example_file <- system.file("extdata/gwaspoly_res_thr.rda",
                                     package = "hidecan")
```

```
gwaspoly_res_thr <- readRDS(gwaspoly_example_file)
```

#### Constructing the HIDECAN plot

The output of the `set.threshold()` function from the `GWASpoly` package can be used as an input for the `hidecan` function `hidecan_plot_from_gwaspoly()`, in order to create a HIDECAN plot representing the GWAS results for each combination of trait and genetic model. The significance thresholds are directly read from the `GWASpoly` output object and do not need to be manually set by the user. In order to improve the clarity of the figure, the argument `remove_empty_chrom` is set to `TRUE`, in order to remove from the plot chromosomes with no significant markers:

```
hidecan_plot_from_gwaspoly(
  gwaspoly_res_thr,
  remove_empty_chrom = TRUE
)
```

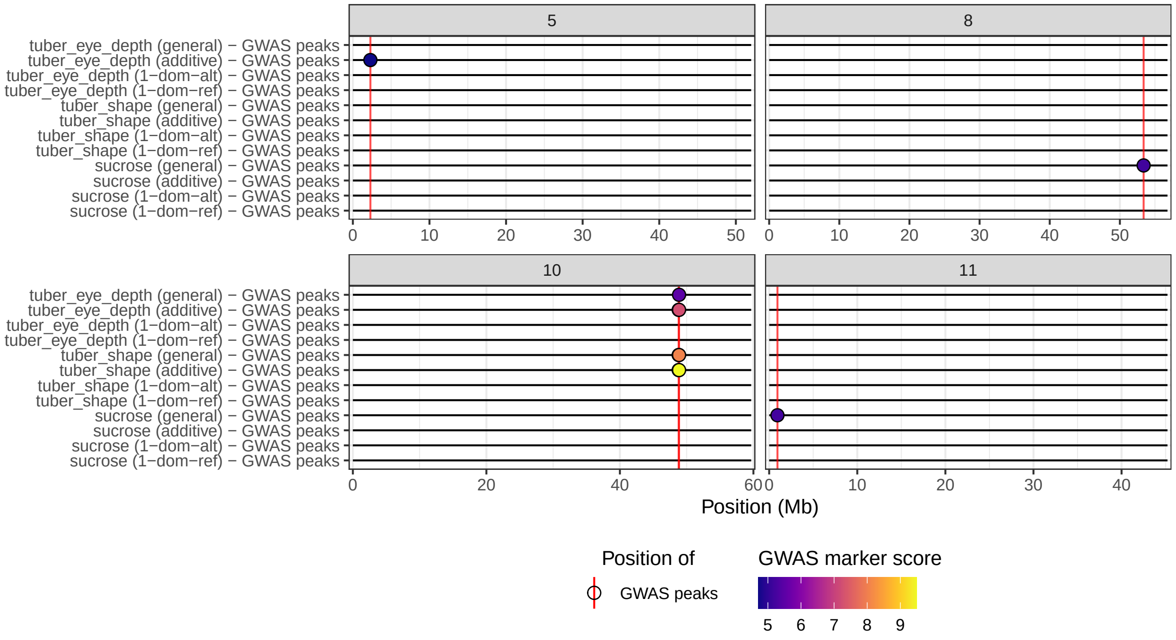


This example is also available online at https://plantandfoodresearch.github.io/hidecan/articles/web_only/gwaspoly_output.html.

### References

Angelin-Bonnet, Olivia, Susan Thomson, Matthieu Vignes, Patrick J. Biggs, Katrina Monaghan, Rebecca Bloomer, Kathryn Wright, and Samantha Baldwin. 2023. "Investigating the genetic components of tuber bruising in a breeding population of tetraploid potatoes." *BMC Plant Biology*.

Rosyara, Umesh R., Walter S. De Jong, David S. Douches, and Jeffrey B. Endelman. 2016. "Software for Genome-Wide Association Studies in Autopolyploids and Its Application to Potato." *The Plant Genome* 9 (2): plantgenome2015.08.0073. https://doi.org/10.3835/plantgenome2015.08.0073.
